# Arabidopsis mutants representing each of the four Mediator modules reveal unique functions in the transcriptional response to salt stress

**DOI:** 10.1101/2022.06.24.497547

**Authors:** Fazelaat Karamat, Alexander Vergara, Jeanette Blomberg, Tim Crawford, Nóra Lehotai, Matilda Rentoft, Åsa Strand, Stefan Björklund

## Abstract

Changes in growth environment trigger stress responses in most organisms. The mechanisms mediating these responses are only partly understood and involve signaling pathways and transcription factors. Mediator is a conserved co-regulator complex required for transcriptional regulation of all eukaryotic protein-encoding genes. However, its function in abiotic stress responses is elusive. We here describe global gene expression changes triggered by salt stress in Arabidopsis. To explore the involvement of Mediator in salt stress response we characterized *med9*, *med16*, *med18*, and *cdk8* mutants representing each of the four modules of Mediator. Our transcriptome data revealed enrichment of shared and specific cis-elements corresponding to unique transcription factors in promoters of mis-regulated genes for each mutant. We show that individual Mediator subunits interact with specific transcription factors to generate a transcriptional stress response and that the mutant phenotypes support the transcriptome data. *med16* and *med18*, and to some extent *cdk8*, display defects in abscisic acid and anthocyanin metabolism and we identify signal molecules, transcription factors and target genes involved in these pathways as dysregulated in the Mediator mutants. Our results reveal how signals from different stress response pathways are dependent on and integrated by Mediator subunits to coordinate a functional response to salt stress.

## Introduction

As sessile organisms, plants have developed advanced ways to adapt to harsh conditions by adjusting their biochemical and molecular properties and they rely on integration of stress-adaptive metabolic and structural changes into endogenous programs. This integration is primarily controlled at the transcriptional level. Salinity is an important plant stressor which has a large impact on plant development and productivity (Flowers, 2004; Prosekov and Ivanova, 2018; Tester, 2010; Agarwal et al., 2013). High salt concentrations in soil decrease the osmotic potential of the soil solution and reduces the accessibility of water around the plant roots. This affects uptake of essential nutrients and reduce their availability for plant cells. Increased Na^+^ and Cl^-^ concentrations in plants induce toxicity, and cause changes in metabolism and hormone levels (Prerostova et al., 2017). These effects together with oxidative damage and impaired photosynthesis can severely restrict plant growth and development.

When plants perceive salinity, signals are conveyed to the nucleus via complex signalling networks (Ulm et al., 2002). This leads to activation of transcriptional programs, which results in activation or repression of stress response genes which subsequently leads to changes in metabolic and physiological activity. Plants use numerous mechanisms to adapt to salinity, such as phytohormone signalling pathways, salt stress responsive protein synthesis (Chinnusamy et al., 2005) , the salt overly sensitive (SOS) pathway (Zhu, 2001; Zhu et al., 1998), transcriptional regulation pathways and cytoskeleton remodelling (Wang et al., 2007a, 2011). The regulatory processes of plant salt stress responses involve control of water flux and cellular osmotic adjustment through synthesis of osmoprotectants (Flowers, 2004; Agarwal et al., 2013; Munns, 2005; Ashraf and Akram, 2009; Yokoi et al., 2002). Increased salt concentrations also lead to negative effects on the redox homeostasis and the cellular energy supply. This is balanced by global re-programming of primary metabolism and by altered cellular architecture (Chen et al., 2005; Baena-González et al., 2007; Jaspers and Kangasjärvi, 2010; Miller et al., 2010; Zhu et al., 2010). Even though several transcriptional regulators that respond to salt stress have been described, the molecular mechanisms involved in salt responses are not fully understood.

Regulation of RNA polymerase II (Pol II) transcription is controlled by chromatin remodelling complexes that affect chromatin structure (Cairns, 2009), by factors that modify histone tails (Wang et al., 2007b), and by transcriptional regulators (activators and repressors) that bind to specific promoter sequences. However, promoter-bound transcriptional regulators are dependent on co-regulators to integrate and to transfer their signals to the general Pol II transcription machinery. Mediator is the most important and well-studied transcriptional co-regulator complex and is required to convey signals from activators/repressors to the Pol II transcription machinery at essentially all gene promoters in eukaryotic cells (Boise et al., 1993; Kim et al., 1994). Mediator subunits have been identified in several eukaryotes from yeast to plants and mammalian cells, and mutations in the corresponding genes have revealed how these subunits function and interact. A combination of genetics, biochemistry and structure biology have shown that Mediator is composed of three modules; Head, Middle and Tail (Dotson et al., 2000; Kang et al., 2001). In addition, a fourth more loosely associated, regulatory Kinase module has also been identified (Liao et al., 1995). Tail subunits are thought to be the main receivers of signals from promoter-bound transcriptional regulators by making direct interactions with activators and repressors, while the Middle module is believed to transfer signals from the Tail to the Head module which in turn makes direct contact with Pol II. The function of Mediator is not merely to transfer information from activators/repressors to Pol II, but also to integrate regulatory information from all transcriptional regulators binding to a promoter at a specific time point, to transcribe each gene at an appropriate level. Plant and human cells comprise more Mediator subunits than yeast, but the overall structure is conserved (Kuang-Lei Tsai, Chieri Tomomori-Sato, Shigeo Sato, Ronald C. Conaway, Joan W. Conaway, 2014).

Genetic analyses have shown that Mediator subunits are involved in different abiotic stress-response pathways in Arabidopsis. Similar to the observations from Arabidopsis, yeast mutations in several Mediator subunits affect the response to different types of stress and in humans Mediator function is connected to multiple diseases (Hashimoto et al., 2011; Huang et al., 2012) . Thus, Mediator plays an essential role in the response to a wide range of stresses. In some cases, the function of the specific subunit has been linked to specific transcription regulators. Med2, Med14 and Med16 have all been identified as important for a proper CBF-dependent response to cold stress (Hemsley et al., 2014). Med8 is involved in response to H_2_O_2_ and *med8* mutants display increased tolerance to oxidative stress (He et al., 2021). Med18 has been linked to abiotic stress through direct interaction with NUCLEOPORIN85 (NUP85), and it was reported that *nup85* and *med18* display hypersensitivity to abscisic acid (ABA) and salt stress as well as overlapping defects in expression of specific stress target genes (Zhu et al., 2017). Furthermore, Med18 has been coupled to ABA signaling through interaction with ABI4 and YY1 in regulation of important abiotic stress response genes (Lai et al., 2014; Li et al., 2016). Finally, Med25 was shown to be important for several stress-response signaling pathways in plants through interactions with multiple transcription factors (TFs) (Blomberg et al., 2012; Chen et al., 2012). Med25 has also been reported to connect the jasmonic acid (JA) receptor COI1 with chromatin and Pol II via the MYC2 TF, and *med25* is sensitive to salt stress but resistant to drought (An et al., 2017; Liu et al., 2019; Elfving et al., 2011). Despite these examples, which indicate that Mediator subunits are involved in multiple signaling pathways, the mechanisms behind the specific responses of the individual Mediator subunits in response to abiotic stress are still elusive.

Here, we use RNA sequencing (RNA-Seq) to identify differentially expressed genes (DEGs) in response to salt stress in wild type (Col-0) Arabidopsis plants and we identify which of these DEGs that are non-responsive (NR) in *med9*, *med16*, *med18*, and *cdk8* (also called *cdke1*) mutants representing each module of the Mediator complex. We also identify transcription factor binding sites (TFBSs) that are enriched in the Col-0 DEG promoters and those that are enriched in the NR promoters of each mutant. Our results suggest key roles for specific subunits of Mediator in integration of signaling pathways induced by salt stress and provide a systems-level evaluation of the regulation of salt stress-responsive transcription by the plant Mediator.

## Results

### The med9, med16, med18 and cdk8 Mediator mutants display unique phenotypes in response to salt stress

To evaluate the function of Arabidopsis Mediator subunits in response to salt stress, we selected mutant lines representing each of the four Mediator modules. One-week old seedlings of *med9* (Middle module)*, med16* (Tail module)*, med18* (Head module), and *cdk8* (Kinase module) were exposed to 200 mM NaCl for two weeks and the number of bleached leaves were quantified. The *med18* mutant showed a significantly higher number of bleached leaves compared to Col-0, indicating that *med18* is sensitive to salt stress (Fig. 1A, 1B). In contrast, fewer bleached leaves were observed for the other mutant lines. In mature plants (5 weeks old) all mutant lines, but especially *med16* and *med18*, exhibited a more sensitive phenotype to salt stress relative to Col-0 when irrigated with a 300 mM NaCl solution for one week (Fig. 1C). The salt sensitivity of all lines was quantified in terms of percent ion leakage from leaves at 72 hours after stress induction (Fig. 1D). Ion leakage was significantly higher for all mutants compared to Col-0 both before and at 72 h after application of stress. For the mutants displaying the strongest phenotypes, *med16* and *med18*, we also made complemented lines (*med16-C* and *med18-C*) expressing HA-tagged versions of Med16 and Med18 under control of their native promoters (Fig. S1A). We found that expression of the HA-tagged version of Med18 could complement the bleaching phenotype of leaves in *med18* (Fig. S1B and C). The *med16-*C also showed complementation of phenotypes observed in *med16* (see below).

**Figure 1.**
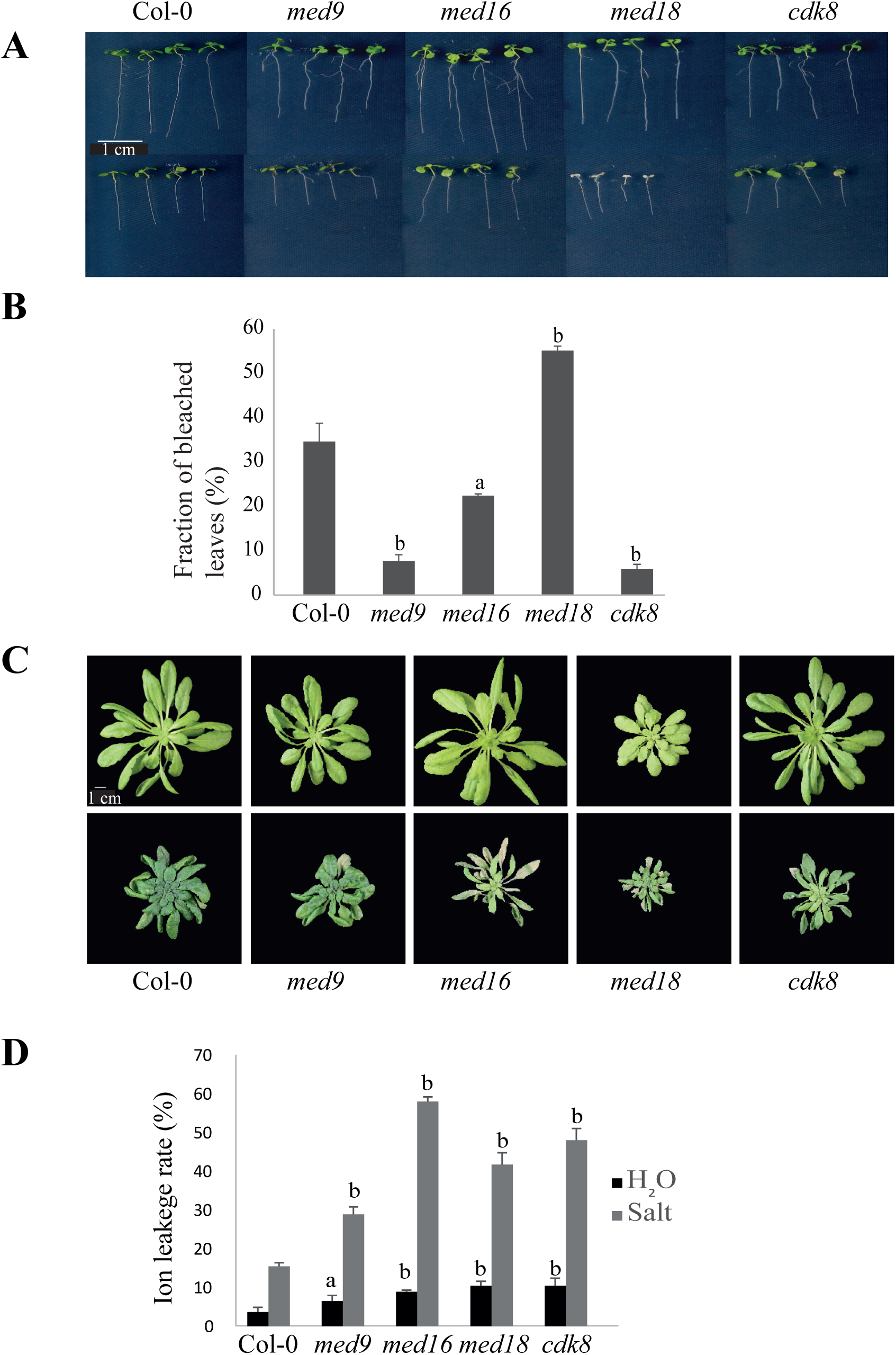
Phenotypic evaluation of Mediator mutants to salt stress. **(A)** Representative images of leaves of Col-0 and each mutant before (upper panel) and after (lower panel) treatment with NaCl. Seedlings were grown for five days at ½ x MS media and then transferred to either ½ x MS media or ½ x MS + NaCl (200 mM). Pictures were taken seven days after transfer. **(B)** Quantification of bleached leaves in (A). Three independent experiments were performed using 20 plants per experiment. **(C)** Representative images of five-weeks-old Col-0 and each mutant irrigated with water (upper panel) or a 300 mM NaCl solution (lower panel). Representative images were taken 7 days after salt stress. (D) Quantification of ion leakage in Col-0 and mutant plants. Ion leakage rates were measured for water irrigated (H2O) and 300 mM salt solution irrigated plants. Three independent experiments were performed using six plants per experiment. Error bars indicate ± SD (n = 3). Statistical analyses were performed comparing Col-0 with each mutant separately. p≤0.05=a, p≤0.01=b (Student’s t-test).

Further analysis of Fig. 1C revealed that leaves of mature *med16* and *med18* became yellow/bleached in response to salt stress indicating loss or degradation of chlorophyll. This phenotype was reversed in the complemented lines (Fig. 2A). To quantify the chlorophyll loss, healthy leaves of Col-0, *med16*, *med18*, and the corresponding complemented lines (*med16-*C and *med18-*C) were incubated for two days in water or in a 200 mM NaCl solution. *med16* and *med18* displayed a significantly higher reduction in chlorophyll content relative to Col-0 and this phenotype was partially reversed in the complemented lines (Fig. 2B). We also found that leaves of *med16* were almost completely degraded after four days of salt stress (Fig. S2A).

**Figure 2.**
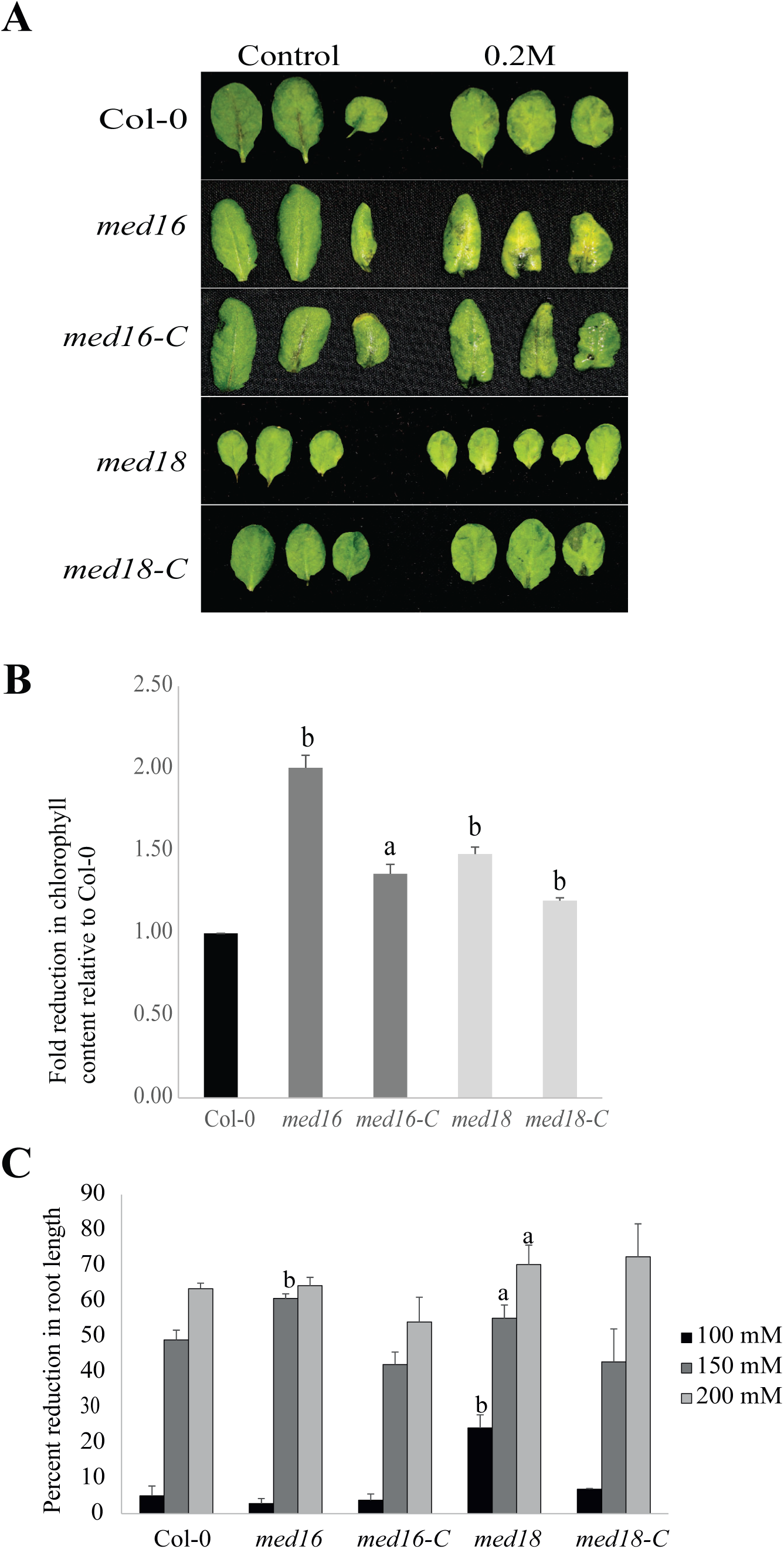
Salt stress affects bleaching, chlorophyll contents, and root growth in Mediator mutants. (**A**) Images of leaf bleaching for Col-0, *med16*, *med18* and the corresponding complemented lines. Leaves taken from five-week-old plants were incubated in 200 mM NaCl and used for phenotypic evaluation. (**B**) Quantification of chlorophyll content in leaves incubated in 200 mM NaCl. Three independent experiments were performed using leaves from 6 plants per experiment. Leaves were subdivided into three replicates. For statistical purposes, the chlorophyll reduction of Col-0 was set to 1. (**C**) Percent reduction in root length of Col-0, *med16*, *med18*, and the corresponding complemented lines when grown at the indicated salt concentrations. Seedlings were grown for one week under normal conditions and transferred to new media without or with the indicated concentrations of NaCl. Measurement of root lengths for each genotype was performed two weeks after transfer using the Image J software. Three independent experiments were performed using 10 plants per experiment. For statistical analysis, the percent reduction was calculated relative to Col-0. Error bars indicate ± SD (n = 3). Statistical analyses were performed comparing Col-0 with each mutant separately. p≤0.05=a, p≤0.01=b (Student’s t-test).

Primary root length is another parameter known to be affected by salt stress in Arabidopsis (Xu et al., 2010). Measurements at different concentrations of NaCl showed that the primary root lengths of *med9* and *med18* were significantly reduced relative to Col-0 already at 100 mM NaCl (Fig. S2B). At 150 mM, also *med16* displayed reduced root length. The root length phenotypes for *med16* and *med18* were supressed in their corresponding complemented lines (Fig. 2C).

#### Global transcriptional responses of wild type Arabidopsis to salt stress

To better understand Mediator function in transcriptional responses to salt stress, we performed RNA-Seq experiments where samples were collected prior to, and at two time points following the application of stress. To achieve an even distribution of salt, all plants were grown hydroponically until reaching mature stage (5 weeks). Salt stress was then imposed by replacing the normal growth media with new media, with or without 200 mM NaCl. RNA was isolated at an early (ET: 4 hour) and a late (LT: 24 hour) time point after salt stress (Fig. S3A). These conditions and time points were selected from pilot experiments aiming to find conditions to avoid dramatic phenotypic effects in any of the mutants (Fig. S3B).

We first analysed the global response to salt stress in Col-0. By selecting genes that showed a log_2_-fold change in expression >0.5 for induced genes and <-0.5 for repressed genes, and an FDR<0.05 at the ET and LT relative to unstressed cells, we identified 5,474 (2,765 induced and 2,709 repressed) and 13,266 (6,734 induced/6,532 repressed) differentially expressed genes (DEGs) at the ET and LT, respectively (Fig. 3A and Table S2). Gene Ontology (GO) analysis showed that each of the four gene lists resulted in enrichment of unique GO categories, where the largest overlap was found between the induced DEGs at the ET and LT (Fig. 3B). The majority of induced DEGs at the ET and LT encodes proteins involved in different types of stress responses (cold, ABA, hypoxia, water deprivation, salicylic acid (SA), salt, wounding etc), transcriptional regulation, autophagy, protein ubiquitination and protein dephosphorylation. Some GOs were unique for the induced DEGs at each time point. Regulation of stomatal closure and response to chitin, defense and wounding were enriched GOs at the ET, while protein modification and transport, leaf senescence and endocytosis were enriched GOs at the LT. GOs enriched among the repressed DEGs were clearly distinct from the induced DEGs. The most highly enriched common GO categories for the repressed DEGs in ET and LT were related to chloroplast function, photosynthesis, glucosinolate biosynthesis and ribosome function/rRNA processing. For the repressed DEGs at the ET, we identified auxin−activated signaling pathway, response to karrikin/red light, xanthophyll biosynthesis, and water transport as unique GOs, while the LT repressed DEGs were exclusively enriched for GOs related to metabolism (methylation, RNA modification, microtubule movement, photorespiration, DNA replication, photosynthesis etc).

**Figure 3.**
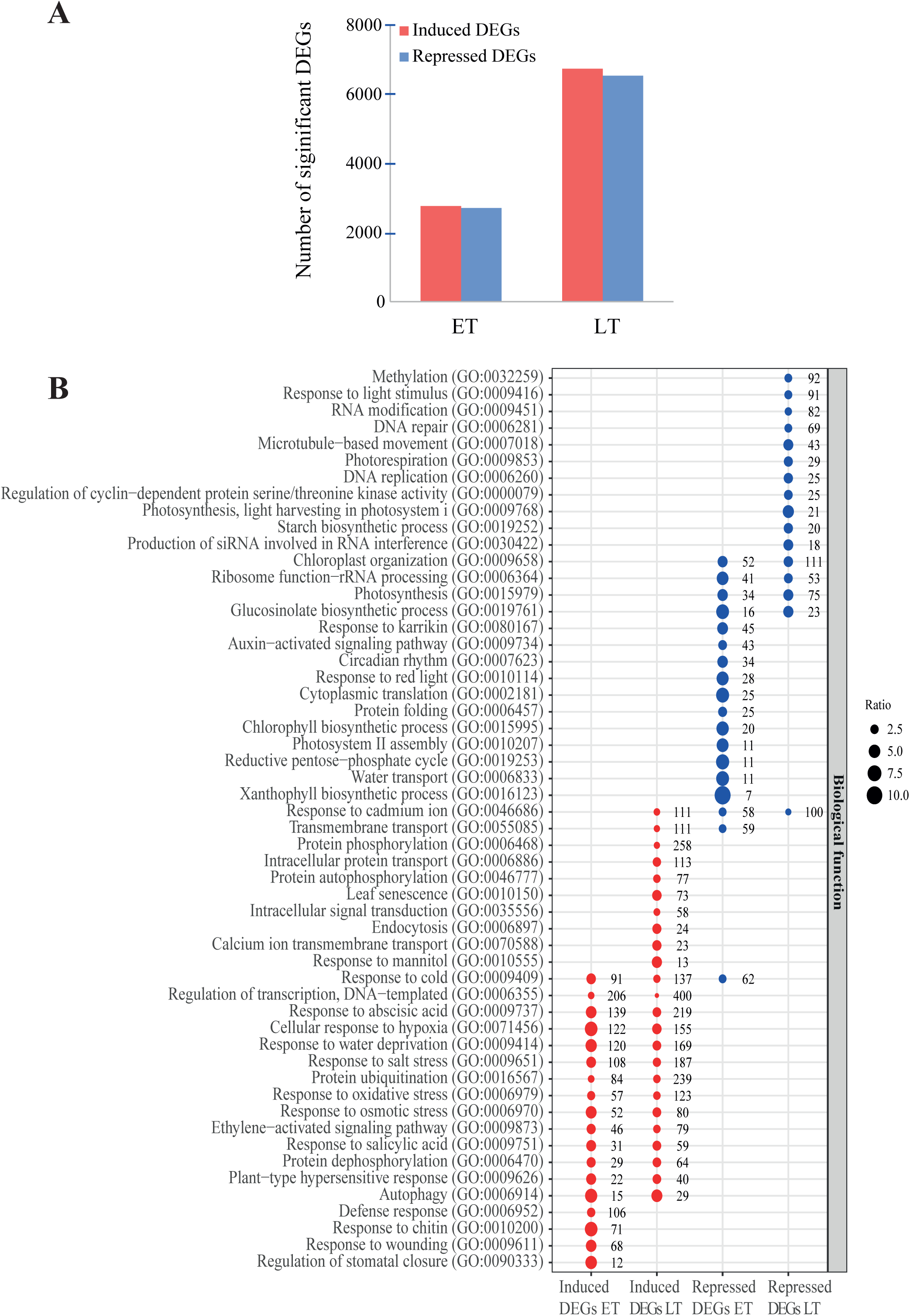
Transcriptional response of Col-0 to salt stress. (**A**) Quantification of induced and repressed DEGs at the Early (ET) and Late (LT) time-points. (**B**) Gene ontology (GO) enrichment analysis for DEGs of Col-0. Significant GOs (p-value ≤ 0.05; Bonferroni adjusted). REVIGO was used to include daughter GO categories to main functional categories. Redundant functional GO categories were removed for the purpose of clarity. Sizes of dots represent the ratio of the abundance of a specific GO category in our experiments relative to the expected ratio in the background data set for each group of genes. Numbers to the right of each dot indicate the number of genes in our data for each functional category.

A main function of Mediator is to interact with promoter-bound TFs and to transmit their signals to the general transcription machinery. We therefore used TF2Network to identify TFBSs enriched in promoters of DEGs that respond to salt stress in Col-0 (Kulkarni et al., 2018). With an aim to focus on the primary transcriptional response to avoid indirect effects, we chose to only search for TFBSs enriched in promoters of the induced and repressed DEGs at the ET. These TFs were grouped into families using the classification defined by the Plant Transcription Factor Database 5.0 (Jin et al., 2017). For the induced DEGs at the ET, we identified 1,066 significantly (p<0.05) enriched TFBSs specific for 642 unique TFs (Table S3). The most common TF-families, each representing >5% of the identified TFs were the ERF-, WRKY-, bHLH-, Myb-, bZIP, and NAC-families (Table S4). Of the 50 most highly enriched individual TFs (=117 TFBSs), 19 (=38%) belonged to the bZIP-family and 18 (=36%) to the bHLH-family (Table S3). In line with our GO analysis, the 10 most highly enriched TFs (ABI5, ABF1, ABF2, ABF4, GBF4, bZIP13, bZIP16, HYH, HY5/bZIP56, and BZR1) are all involved in different aspects of ABA-responses (Bensmihen et al., 2002; Choi et al., 2000; Hsieh et al., 2012; Yang et al., 2018, 2016). Seven of these TFs were by themselves induced at the ET (Table S5), suggesting that they function as transcriptional activators important for the early response to salt stress. We also noticed a high proportion of six TFs belonging to a specific group of Myb-related TFs among the top 50 TFs. EPR1/RVE7, LCL5/RVE8, LCL1/RVE4, RVE6, LHY, and RVE5 (Table S3) all belong to a sub-family of the Myb-related TFs called the CCA/LHY/RVE gene family that have important functions in a molecular network that controls the Arabidopsis circadian clock (Gray et al., 2017). Different from the bZIP-family TFs, expression of all, except two, of the CCA/LHY/RVE were repressed at the ET relative to the control (Table S5), suggesting that they function as transcriptional repressors in the early response to salt stress. In line with this, TFBSs for the seven bZIP TFs and six CCA/LHY/RVE TFs were all significantly enriched in promoters of DEGs that were induced at the LT. This indicates that TFs driving the early transcriptional response to salt stress also are important for the late transcriptional response (Table S6).

In promoters of repressed DEGs at the ET, we identified 528 significantly enriched TFBSs, corresponding to 384 unique TFs (Table S7). The most common TF-families, each representing >5% of the identified TFs were the ERF-, Myb-, bZIP-, HD-ZIP-, MIKC/MADS-, and Dof- families (Table S8). We chose to exclude TFs that have not yet been classified as part of any TF-family (labelled N.D in Table S7). The three first families were enriched among the induced DEGs at the ET as well, while the HD-ZIP, MIKC/MADS and Dof-families were specific for the repressed DEGs at ET. Of the fifty most enriched individual TFs (=66 TFBSs) in the repressed DEG promoters at the ET, ten belonged to the TCP-family, nine to the Myb-family, and eight were AGAMOUS-LIKE (AGL)-TFs belonging to the MIKC/MADS-families of TFs (Table S7). When comparing the expression levels of the individual TFs of these families before and at the ET after salt stress, we found that the expression of TCPs in general was repressed (Table S9). In particular, the expression levels of TCP5, 6, and 23 were all significantly repressed, while the expression levels of the Myb-family TFs (Myb10, 32, 43, 51, 99, At2g38090, and At5g08520) were significantly induced. Of the MIKC/MADS TFs, two (AGL68 and PI) showed a significant induction at the ET relative to the control. Our results therefore suggest that TCP TFs before stress typically function as activators for DEGs that are repressed in response to salt stress. In contrast, Myb and MIKC/MADS TFs seem to function as repressors for DEGs that are repressed in response to salt stress. Accordingly, nearly all TFBSs for the TCP, Myb and MIKC-MADS proteins identified here were identified as enriched in promoters of repressed DEGs at the LT after salt stress (Table S10).

#### Col-0 DEGs at the ET can be divided in five clusters controlled by unique transcriptional regulatory programs

We performed heat map analysis based on Z-scores of the 5,474 induced and repressed DEGs in Col-0 at the ET (Table S2) and identified five clusters, each comprising genes that exhibit similar patterns of transcriptional regulatory response at the ET and LT after salt stress (Fig. 4A). Clusters 1-5 were composed of 629, 1,701, 453, 611 and 2,080 DEGs, respectively (Table S11). We performed GO (Fig. 4B) and TFBS analysis (Fig 4C) for each of these clusters.

**Figure 4.**
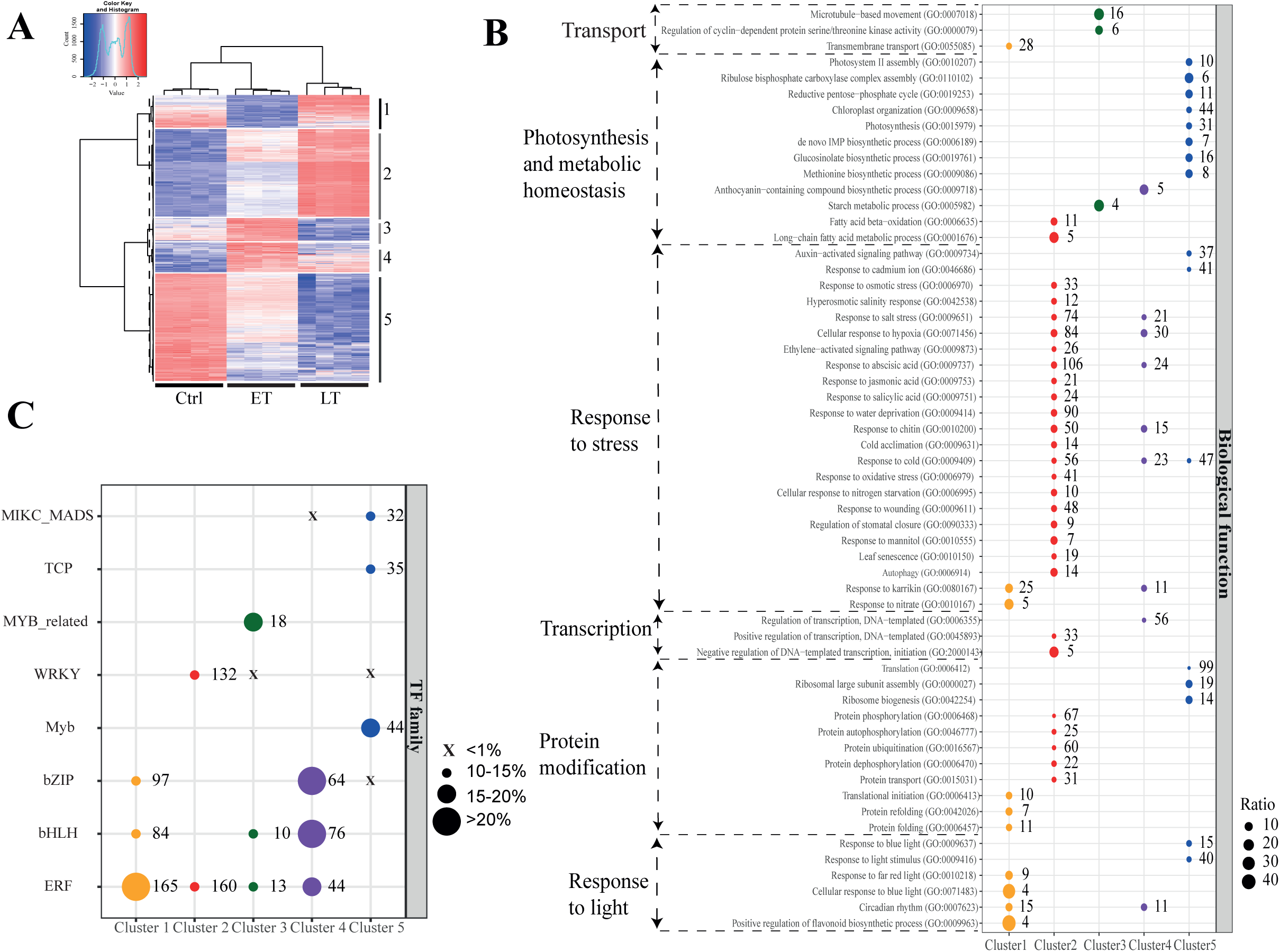
Hierarchical clustering of genes in Col-0 that respond to salt stress. (**A**) Hierarchical clustering of co-expressed genes which are differentially regulated in response to salt stress. VST normalised data of the 5,474 ET DEGs are displayed as z-scores, and cluster dendrograms are shown with a dashed line indicating divisions between 5 co-expressed clusters. Relative expression levels of the ET DEGs are shown for the Control (Ctrl), early (ET) and late (LT) time-points for the four biological replicates. (**B)** Gene ontology (GO) enrichment analysis for DEGs in each cluster. Significant GOs with a Bonferroni adjusted p-value ≤ 0.05 were extracted. REVIGO was used to include daughter GO categories into main functional categories. Redundant functional categories were removed for clarity purposes. Sizes of dots indicate the ratio of the abundance of a specific category in our data compared to expected ratio in background data set for each group of genes. (**C**) Enriched and absent TFBS-families for each co-expressed cluster of early DEG promoters in Col-0. Numbers to the right of dots (in B and C) indicate the number of genes and TFBSs, respectively, present in our data for each category.

We first analysed Clusters 2-4, which are composed of DEGs that are induced at the ET. They split into three clusters that differ in that Cluster 2 DEGs show low expression levels before stress, and then continuously increased levels at the ET and LT. Cluster 3 DEGs are induced at the ET, and then repressed to levels below control (before stress) at the LT. Finally, Cluster 4 DEGs show a similar pattern as Cluster 3 DEGs but are at the LT repressed to levels above their levels in unstressed cells. GO analysis revealed distinct patterns for each cluster indicating that they encode proteins involved in unique salt stress response pathways (Fig. 4B). Cluster 2 was enriched for genes encoding proteins involved in a wide range of stress responses, transcription, and protein modification/metabolism. Cluster 3 was enriched for proteins involved in cell cycle and starch metabolism, while Cluster 4 was enriched for proteins involved in different types of stress responses, transcriptional regulation, circadian rhythm, and anthocyanin biosynthesis. In line with this, analysis of enriched TFBSs (Table S12) and the TF-families the corresponding TFs belong to (Table S13; Fig. 4C), indicated that expression of genes in these three clusters are controlled by distinct transcriptional programs in response to salt stress. The most common TF-families (>10%) in promoters of genes in Cluster 2 belonged to the WRKY- and ERF-families which constitute 12 and 15%, respectively, of the identified TFBSs in this cluster. WRKY and ERF TFs have previously been identified in different types of stress responses in Arabidopsis (Banerjee and Roychoudhury, 2015; Xie et al., 2019). The most enriched TF-families in Cluster 3 belonged to the Myb-related-, ERF- and bHLH-families. We note that all the Myb-related TFs we identify here belong to the CCA/LHY/RVE gene family described above. As seen in Table S12, their TFBSs constitute 19 of the 20 most highly enriched in Cluster 3. Myb-related, ERF- and bHLH TFs have all been reported to control stress responses (Li et al., 2019; Guo et al., 2021a; Xie et al., 2019). Finally, Cluster 4 DEG promoters were enriched for the bHLH-, bZIP-, and ERF-families which have all been implied in abiotic stress (Banerjee and Roychoudhury, 2017; Guo et al., 2021a; Xie et al., 2019). Combined with our results showing that DEGs in different clusters are enriched for different GO categories, the unique enrichment of TFBSs in each cluster provides further support to the model where each cluster is regulated by unique transcriptional programs to control distinct salt stress response pathways.

Clusters 1 and 5 include DEGs that are repressed at the ET after salt stress. However, while Cluster 1 DEGs were induced to levels above those of unstressed cells at the LT, DEGs in Cluster 5 were even further repressed at the LT (Fig. 4A). GO analysis revealed that DEGs in Cluster 1 are involved in circadian rhythm, flavonoid metabolism, transmembrane transport, and responses to light, karrikin and nitrate (Fig. 4B). Interestingly, the GO category “Response to karrikin” was also enriched among the DEGs in Cluster 4 (Fig. 4B), which indicates that expression of genes involved in response to karrikin are both induced and repressed by salt stress. DEGs in Cluster 5 encode proteins involved in different processes related to protein synthesis, metabolism, and photosynthesis, which is in line with the established knowledge that plants adapt to stress by slowing down metabolism (Zhu, 2002). Analysis of enriched TFBSs (Table S12) and TF-families (Table S13) showed that Cluster 1 DEG promoters were enriched in binding sites for the ERF-, bZIP- and bHLH-families, while Cluster 5 was enriched for the Myb-, TCP-, and MIKC/MADS- families. As for the enriched TF-families in Clusters 2-4, the Myb-, and TCP-, the MIKC/MADS TF-families enriched in Clusters 1 and 5 have been linked to different types of abiotic stress responses (Castelán-Muñoz et al., 2019; Dubos et al., 2010; Li, 2015).

We noticed that Clusters 1 and 4 were enriched in the same three TF-families and that TFs from the ERF-family were enriched in all clusters, except Cluster 5. This was surprising since Cluster 1 is composed of DEGs that are repressed, while Clusters 2-4 comprise DEGs that are induced in the early response to salt stress (Fig. 4C). To analyse these results at the level of individual TFs, we made lists and Venn diagrams illustrating the overlaps of the ERF-family position weight matrices (PWMs = TFBSs) identified in each of Clusters 1-4 (Table S14; Fig. S4). Interestingly, we found a large overlap between ERF TFBSs in promoters of DEGs in Clusters 2-4, which comprise genes that are induced by salt stress, relative to the promoters of the represses DEGs in Cluster 1. All ERF TFBSs identified in Clusters 3 and 4 were also found as ERF TFBSs in Cluster 2 (Figs. S4A and S4DD). In contrast, the ERF TFBSs in Clusters 3 and 4 showed only 0% and 4.5% overlap with Cluster 1 (Figs. S4A and S4C). The overlap between Clusters 1 and 2 was 44% (Figs. S4A and S4B). We conclude that distinct ERF TFs are involved in activation and repression of target genes in response to salt stress.

### Effects of Mediator mutants on the salt stress response

After defining the Col-0 response to salt stress, we focused on how the ET salt-stress induced regulon of Col-0 DEGs is regulated in our Mediator mutants. Mediator is primarily involved in transcriptional regulation. In order to identify salt stress-induced genes that are mis-regulated in the Mediator mutants, we analysed the 5,474 genes that are DEGs at the ET in Col-0 and made lists of those that were non-responsive (NR) to salt stress in each mutant (Table S15). NR genes were defined as those showing a log_2_ ratio (fold change at ET vs control in mutants)/ (fold change at ET vs control in Col-0) >0.5 for repressed and <-0.5 for induced Col-0 DEGs. *med9* had 673 NRs (433 not induced/240 not repressed), *med16* had 1,021 NRs (666 not induced/355 not repressed), *med18* had 2,008 NRs (1,261 not induced/747 not repressed) and *cdk8* had 1,358 NRs (964 not induced/394 not repressed) (Fig. 5A; Table S15). Thus, each mutant showed significant effects on regulation of genes that respond to salt stress in Col-0. The *med9* mutant had the smallest (12% of the Col-0 DEGs), while *med18* had the largest effect and caused mis-regulation of 37% of the ET Col-0 DEGs.

**Figure 5.**
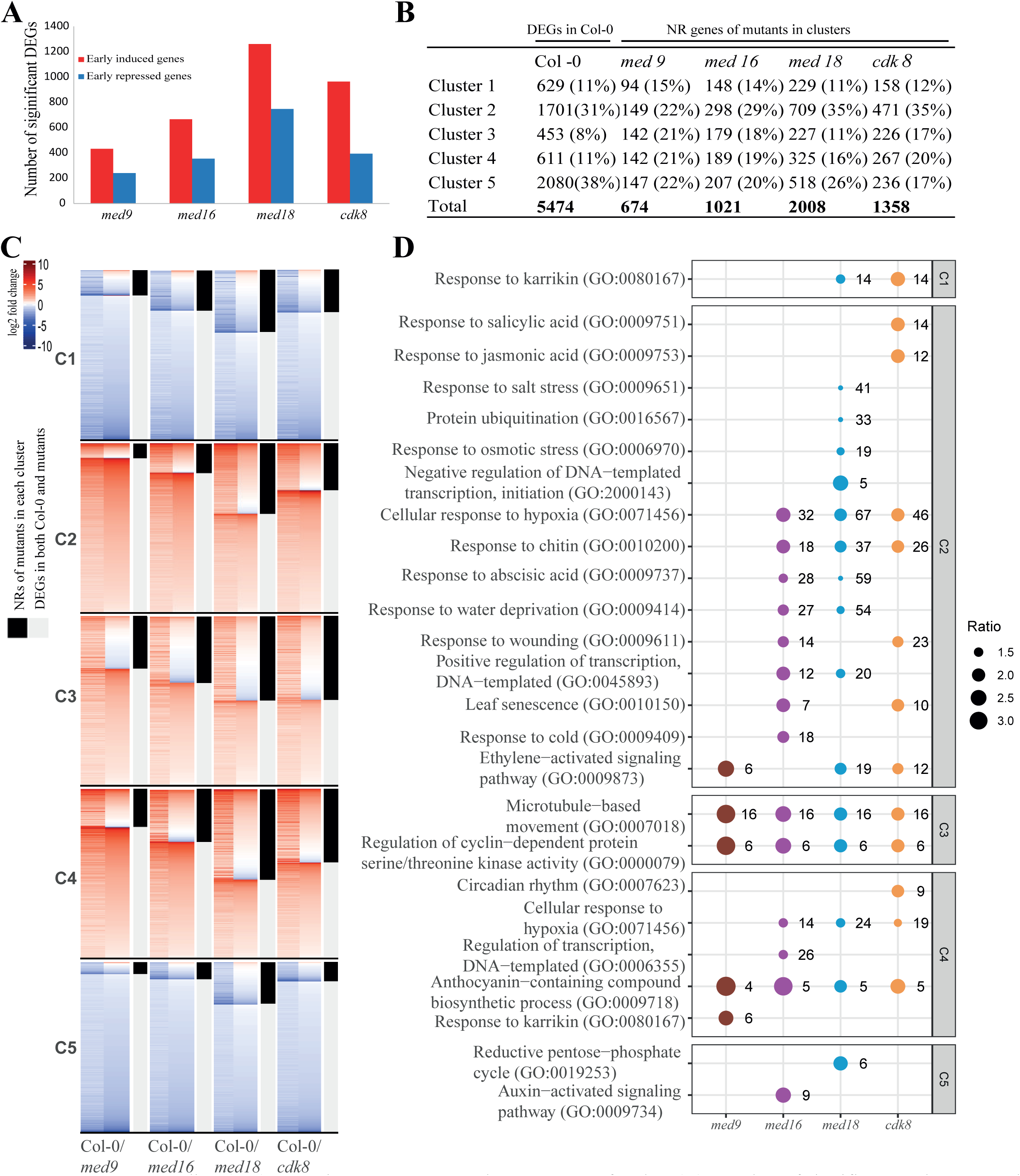
Non-responding (NR) genes in mutants among the ET DEGs of Col-0. (**A**) Number of significant early UPR and DR DEGs of Col-0 that are NR genes in mutants. (**B**) Distribution of NR genes of mutants in each cluster. (**C**) Heatmaps for NR genes in each mutant and cluster. The left column of each heatmap represents fold change in expression of DEGs in Col-0. For each heatmap, left columns represents fold change in expression of DEGs in Col-0. Right columns represent fold increase of the corresponding genes in each mutant. Black cells to the right indicate if a gene is nonresponsive (NR). Gray cells represent genes that are also DEGs in mutants. (**D**) GO enrichment analysis for NR genes in each mutant and cluster. Significant GOs (Bonferroni corrected p-value < 0.05) were determined for each cluster and mutant using the early DEGs of Col-0 in each cluster as background dataset. REVIGO was used to include daughter GO categories into main functional categories. The sizes of dots indicate the ratio of occurrence of a specific functional category for NR genes of mutants versus Col-0. Numbers to the right of dots indicate the number of NR genes of mutants for each functional category.

For analyses of the expression patterns of Col-0 DEGs in the mutants, we made lists of all DEGs in each cluster and mutant and divided them into NR-genes (see definition above) and genes that respond similarly in mutants and Col-0. Genes in these 40 lists (10 for each mutant/8 for each cluster) were sorted separately according to their fold change in the mutants (Table S16). We found that none of the mutants had a major effect on repression of DEGs in Cluster 1, since the number of NR-genes in Cluster 1 relative to the total number of NRs for all clusters (11-15%) was similar in all mutants and did not differ significantly from the number of DEGs in Cluster 1 for Col-0 relative to the total number of DEGs in Col-0 at the ET (11%) (Fig. 5B). In contrast, all mutants were overrepresented for NRs in Cluster 4 (16-21% in mutants vs 11% in Col-0) and underrepresented for NRs in Cluster 5 (17-25%) relative to Col-0 (38%), while *med9*, *med16* and *cdk8* were overrepresented for the number of NRs in Cluster 3 (21%, 18%, 17%, respectively) relative to DEGs in Col-0 and NRs in *med18* (8% and 11%, respectively). Finally, *med9* was underrepresented for NRs in Cluster 2 (22%) relative to Col-0, *med16*, *med18*, and *cdk8* (29-35%). Cdk8 has usually been implicated in negative regulation of transcription. We were therefore surprised to find that *cdk8* was the most underrepresented mutant in Cluster 5, a cluster which is characterized by DEGs that are repressed in Col-0 in the ET response to salt stress.

To further illustrate the effects of Mediator mutants on the ET transcriptional response to salt stress, we used the 40 gene lists in Table S16 to produce heat maps where a red colour represents a higher ratio between the expression level at the ET/the expression level before stress, while a blue colour represents a lower ratio for each line and each cluster (Fig. 5C). For each list, we first sorted genes into those that for each mutant and cluster respond similar to Col-0 (represented by the grey bars to the right of each heat map) and genes that do not respond as Col-0 (NR genes represented by black bars to the right of each heat map). Within each list, genes were sorted such that those with the highest ratio (expression level at ET/expression level before stress) in the mutants are at the top, and those with the lowest ratio are in the bottom of each heat map. In each cluster and mutant, we found that some of the NRs were not only non-responsive but had completely reversed their response to salt stress relative to their response in Col-0 (see red labelled NR genes at the top of each mutant in Clusters 1 and 5, and blue labelled genes at the bottom of the NR genes of each mutant in Clusters 2-4). In general, our analysis again showed correlation to our phenotypic characterization where *med18* displayed the largest, and *med9* the smallest, defects in the ET transcriptional response to salt stress.

GOs analysis of the NR genes for each cluster and mutant showed that NRs in *med9* are uniquely enriched for karrikin genes in Cluster 4, but similarly to the other mutants, enriched for cell-cycle related processes in Cluster 3 and ethylene signalling and anthocyanin processes in Clusters 2 and 4, respectively (Fig. 5D). The *med16*, *med18*, and *cdk8* mutants all showed defects in regulation of different stress response genes in Clusters 2, 4 and 5. However, *med16* was uniquely affected in regulation of cold response genes in Cluster 2, genes involved in transcriptional regulation in Cluster 4 and auxin-signalling genes in Cluster 5. The *med18* mutant was unique in being important for regulation of salt and osmotic stress responses, ubiquitination, and repression of transcription in Cluster 2, and for repression of genes involved in the pentose phosphate cycle in Cluster 5. Finally, Cdk8 was uniquely important for regulation of SA and JA responses in Cluster 2 as well as for regulation of genes involved in circadian rhythm in Cluster 4.

### Analysis of TFBSs enriched in promoters of Cluster 1-5 in each mutant

The results in Fig. 5 indicate that each of the four Mediator subunits are involved in specific transcriptional programs regulating the early transcriptional response to salt stress. We therefore used TF2Network to search for TFBSs in promoters of NRs in each cluster and mutant (Table S17). TF2Network identifies enrichment of TFBSs in a set of promoters relative to their abundance in all promoters of the Arabidopsis genome. However, the lists of NR-genes for each cluster and mutant comprise subpopulations of the DEGs in the corresponding clusters in Col-0. A direct comparison between the lists of TFBSs in Table S17 with those in Table S12 would therefore not identify TFs that are dependent on specific Mediator subunits for the ET response to salt stress since enrichment of TFBS would be biased by the background enrichment of each cluster. To identify TFBSs that are truly enriched in the NR-gene promoters of each mutant and cluster relative to the DEG promoters of Col-0, we used hypergeometric distribution testing (Eden et al., 2007).

#### med9

For *med9*, we identified two TFBSs specific for the GBF3 and TCP24 TFs as significantly enriched in promoters of NR-genes in Cluster 1 relative to the DEG promoters of Col-0 in Cluster 1 (Table S18). This suggests that these TFs are important for repression of target genes in Cluster 1 in response to salt stress and that they depend on Med9 for this function. GBF3 was also the most enriched TF in Cluster 1 of Col-0 (Table S12). It encodes a bZIP G-box binding protein whose expression is induced by ABA and contributes to improved tolerance to osmotic-, salt- and drought-stress (Ramegowda et al., 2017). In contrast, TCP24 was found further down in the list of enriched TFs in Cluster 1 of Col-0 (Table S12) and has not been connected to stress before. However, it has been identified as a transcriptional repressor (Li et al., 2012), which is in line with the repressive response of genes in Cluster 1 and suggests that Med9 is important for repression by TCP24. In Cluster 2, we identified eight significantly enriched TFBSs specific for five different TFs in the promoters of the NR genes in *med9*. Four of these TFs belong to the bZIP-family and one to the Myc-type bHLH-family. The bZIP-family represent 21 of the 50 most enriched TFs in Cluster 2 of Col-0 (Table S12) and they are thus likely to function as important transcriptional activators that require Med9 for induction of genes in response to salt stress. Our results support previous reports showing that these bZIP-family TFs (ABF2, ABF4, ABI5, and GBF2) are involved in ABA signalling and in response to salt stress (Choi et al., 2000; Tian et al., 2020b). Furthermore, the BIM1 bHLH/Myc TF has been linked to brassinosteroid-signaling, which is known to play an essential role in salt stress responses through crosstalk with ABA signalling (Planas-Riverola et al., 2019). For the promoters of Cluster 3 NR-genes in *med9*, we found enrichment of eight TFBSs, each specific for unique TFs that function in Med9-dependent activation of genes in Cluster 3. Five of these belong to the Myb-family (Myb3R-type), two to the homeobox-family and one to the C2H2-family. MYB3R TFs have been identified as repressors which function as key regulators in active growth repression under salt stress, by negatively regulating the transcription of G2/M-specific genes (Okumura et al., 2021). However, neither of RLT1, RLT2 or ZFP8 have previously been associated to stress responses. No TFBSs were significantly enriched in the NR-genes of Cluster 4 in *med9*, and this was true also for the other three mutants (Tables S18-S21). Finally, the NRs-gene promoters in *med9* in Cluster 5 were enriched for nine TFBSs specific for Myc2 and five TCP-family TFs. These TCPs are different from TCP24 that was enriched in Cluster 1, which suggests that specific TCPs can function in unique transcriptional stress-responsive pathways. However, TCPs seem to function in Med9-dependent transcriptional repression in both cases. Overexpression of Myc2 has previously been shown to cause susceptibility toward salt stress (Verma et al., 2020), and some of the identified TCPs have been reported to function as transcriptional repressors in different systems (Hervé et al., 2009; Viola et al., 2016).

#### med16

WRKY17 was the only TF whose TFBS was enriched in the NR-gene promoters of Cluster 1 in *med16* (Table S19). WRKY17 has been reported to be induced by salt and other stresses, and *wrky17* is susceptible to salt stress (Ali et al., 2018). In Cluster 2, we identified three enriched TFBSs specific for two TFs (NAC25 and NAC56) in the *med16* salt- and other types of stress (Tran et al., 2004). The NR-gene promoters of *med16* in Cluster 3 were enriched for essentially the identical set of TFBSs as *med9*, indicating that the corresponding TFs are dependent on both Med9 and Med16 for activation of the salt stress response genes in this cluster. No TFBSs were enriched in the promoters of NR-genes in *med16* in Clusters 4 or 5.

#### med18

Analysis of the NR-gene promoters in Cluster 1 of *med18* revealed one enriched TFBSs specific for WRKY27 (Table S20). Mutations in WRKY27 in rice (*Oryza sativa*) have been shown to be resistant to salt stress (Pang et al., 2017). The highest number of TFBSs for all mutants and clusters were found in the list for *med18* in Cluster 2. This is in line with several previous reports indicating that Med18 is one of the most important non-essential Mediator subunits (Fallath et al., 2017; Lai et al., 2014). In summary, we identified 41 enriched TFBSs representing 29 unique TFs in the promoters of NR genes in Cluster 2 of *med18*. Fifteen of these TFs belong to the bHLH-, 7 to the bZIP-, 4 to the TCP-, 2 to the CAMTA-, and 1 to the ARF-families. Most of these TFs have previously been reported to be involved in different types of stress responses in Arabidopsis. The most notable were NaCL-INDUCED GENE 1 (NIG1), which encodes the first known plant TF involved in direct Ca2+ binding (Kim and Kim, 2006), and the calmodulin-binding transcription activators CAMTA2 and CAMTA3, which have been implicated in different types of stresses (Iqbal et al., 2020). This suggests that Med18 is important to respond to Ca^++^- dependent stress-responses. In Cluster 3, the NR-genes in *med18* were enriched for two TFBSs, each specific for a unique TF, MYB3R1 and MYB3R5. These are members of the Myb3R TF-family which was also enriched in the NR-gene promoters in Cluster 3 of *med9* and *med16*. However, the NR-genes in Cluster 3 of *med18* were not enriched for the TFBSs specific for the HB-other and C2H2 TFs that were identified as enriched in *med9* and *med16*. Finally, six TFBSs representing five unique TFs were enriched in the NR-genes of Cluster 5 in *med18*. Two of them, Myc2 and Tcp19, were also identified in the NR-genes in Cluster 5 for *med9*. However, HDG1, AHDP and HP15 were uniquely enriched in *med18*, and are all TFs of the homeodomain-ZIP TF-family. Both HDG1 and AHDP have been reported as important for formation of the cuticle, a chemically heterogeneous lipophilic layer that protects plants from biotic and abiotic stresses (Nadakuduti et al., 2012; Wu et al., 2011), and HB15 was identified as part of transcriptional regulatory networks in Arabidopsis during single and combined stresses (Barah et al., 2015).

#### cdk8

For *cdk8*, we found no enriched TFBSs in Cluster 1 (Table S21). For Cluster 2 NR-promoters we identified 8 TFBSs specific for 5 unique TFs. Two of these (HY5 and CAMTA3) were also identified in Cluster 2 of *med18*. HY5 has been reported as important for response to salt stress (Yu et al., 2016), ABA (Chen et al., 2008) and in positive regulation of anthocyanin metabolic processes (Zhang et al., 2011). CAMTA3 has been shown to be involved in responses to both salt (Iqbal et al., 2020), and cold stress (Doherty et al., 2009). Interestingly, CAMTA3 is also important for cold-induced expression of the drought- and cold-responsive TFs DREB1A/CBF3, DREB1B/CBF1 and DREB1C/CBF2 (Kidokoro et al., 2017), which all were found as enriched in the NR-promoters of this cluster in *cdk8*. Furthermore, a mutation in *CDK8* was reported to suppress the dwarfism and constitutive cold resistance phenotypes of a *camta1/2/3* mutant (Huang et al., 2019). As described for the other three mutants, the NR-promoters of Cluster 3 in *cdk8* were enriched for a partially overlapping set of Myb TFBSs. Finally, the *cdk8* NR-gene promoters in Cluster 5 were enriched for 6 TFBSs interacting with 4 TFs. Three of these are specific for the TCP family which was also enriched in Cluster 5 of *med9* and *med18*. However, none of the TCP TFs that were enriched in *cdk8* was common to those enriched in the *med9* and *med18* NR-promoters. Notably the three TCPs that were uniquely identified in Cluster 5 of *cdk8*; TCP5, TCP13 and TCP17, have been shown to play fundamental roles in promoting thermoresponsive hypocotyl growth by positively regulating the activity of the PIF4 TF (Zhou et al., 2019). The last identified TF was Myc2, which was also enriched in Cluster 5 of both *med9* and *med18* (see above).

To obtain an overview of the transcriptional salt stress response, we first focused on how genes in each cluster are regulated and how they depend on each of Med9, Med16, Med18, and Cdk8. We therefore summarized enrichment of TFBSs at the level of TF-families for NR genes in each cluster and mutant and compared it to the enrichment of TF-families for each cluster in Col-0 (Table S22; Fig. S5). Cluster 1 NR gene promoters were enriched for bZIP- and TCP-family TFBSs in *med9* and WRKY-family TFBSs in *med16* and *med18*. Cluster 2 NR gene promoters were enriched for bHLH- and bZIP-TFBSs in *med9*, NAC-family TFBSs in *med16*, bHLH-, bZIP-, TCP-, CAMTA- and ARF-families in *med18*, and ERF-, bZIP- and CAMTA-families in *cdk8*. In Cluster 3, Myb-family TFs were enriched in NR gene promoters of all mutants. In addition, the C2H2- and HB-other families were enriched in both *med9* and *med16*. No enriched TF-families were detected in NR gene promoters of Cluster 4 for any of the mutants. Finally, the *med9*, *med18* and *cdk8* NR gene promoters of Cluster 5 were all enriched for TCP- and bHLH- families and *med18* was also enriched for the HD-Zip family.

As an alternative way to visualize the salt stress response, we constructed a gene regulatory network focused on each of the Mediator mutants. Fig. 6 show the upstream TFs and downstream target genes which are dependent on each of the four Mediator subunits for proper function. To make it easier to search for the TFs that depend on each of the Mediator subunits and their corresponding target genes are also presented in the form of a table (Table S23).

**Figure 6.**
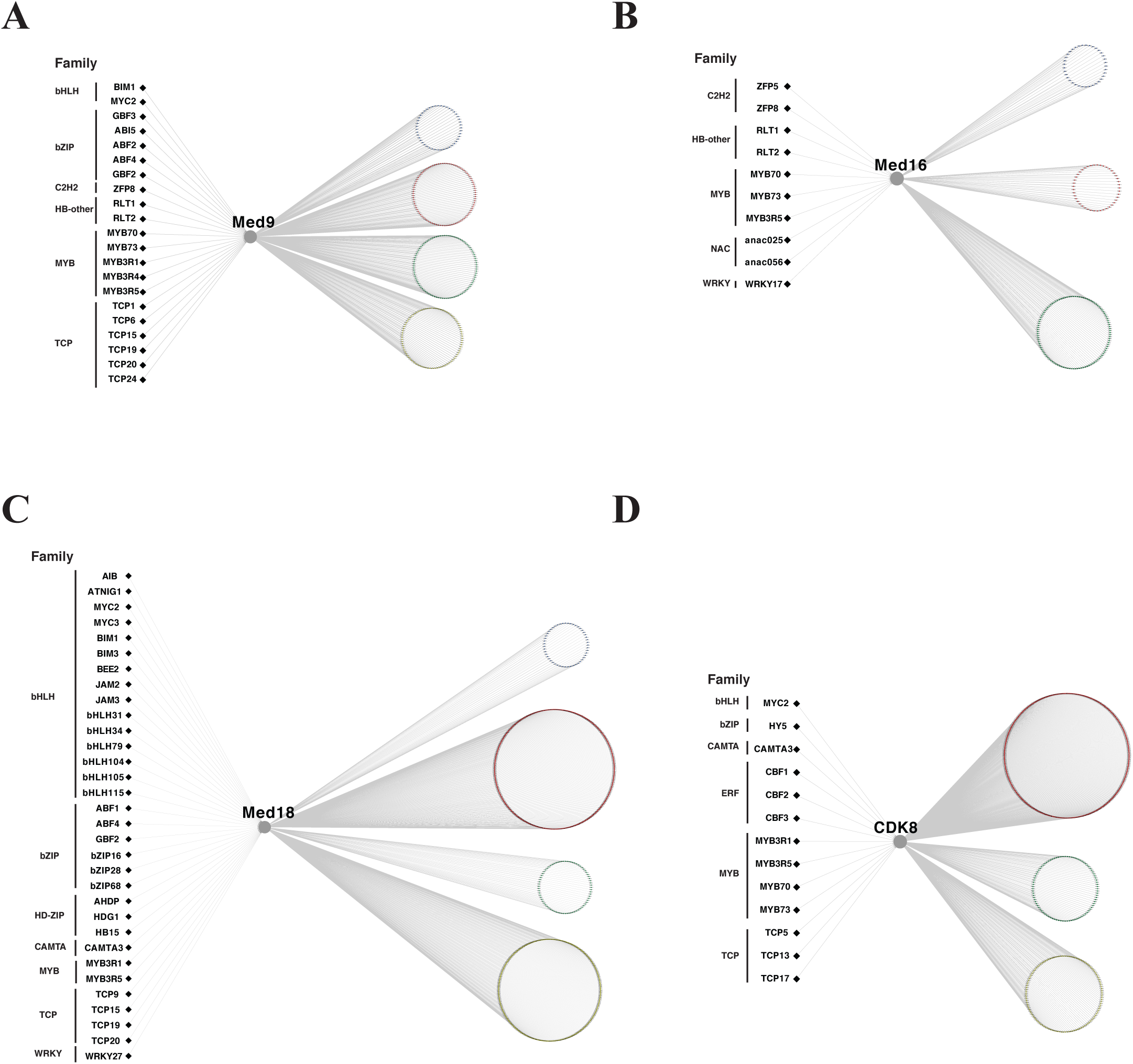
Gene regulatory networks of Mediator subunits. For each analyzed mediator subunit, we built regulatory networks by means of the identification of mis-regulated genes in *med9* (A), *med16* (B), *med18* (C), and *cdk8* (D). Transcriptions factors (black diamonds) and their family memberships are represented to the left. Mediator subunits and target genes are represented by circles. Target genes were colored according to their membership of the five clusters (C) identified in Figure 4A. Members of cluster C1, C2, C3 and C5 are shown in blue, red, green and yellow respectively.

### The med16, med18, and cdk8 mutants show multiple defects in transcriptional pathways that control anthocyanin metabolism and display defects in induction of anthocyanin levels in response to salt stress

Salt stress causes ionic and osmotic stress in plants and thereby reduces their capacity for water uptake and maintenance of photosynthesis. Salt stress also results in nutrient imbalances and damage to cell membranes. Anthocyanins belong to a parental class of molecules called flavonoids, which are synthesized via the phenylpropanoid pathway. Their production is triggered by several abiotic stresses in plants, for example increased salt concentration. Anthocyanins mainly serve as antioxidants which scavenge excessively produced ROS under stress conditions (Ai et al., 2018; Chalker-Scott, 2002; Naing et al., 2018; Wang et al., 2016), and give plants a reddish to purple colouring. In our experiments, we detected purple colouring in Col-0 and all mutants, but it was less pronounced in *med16*, *med18* and *cdk8* (Fig. S6A). This experiment was performed at short day conditions, but we found that the effect on *med16*, *med18* and the corresponding complemented lines was even more pronounced at long day conditions (Fig. S6B).

In agreement with these phenotypic results, we found that the GO category “Anthocyanin−containing compound biosynthetic process” (GO:0009718) was enriched among the NR genes in Cluster 4 of all four Mediator mutants (Fig. 5D). Four genes involved in anthocyanin metabolism (*AT4G14090*, *LDOX*, *DFR*, and *UF3GT*) were identified as NRs in Cluster 4 of all four mutants (Fig. 7A upper panel; Table S24). In addition, *med16*, *med18* and *cdk8* showed defects in induction of the *PRODUCTION OF ANTHOCYANIN PIGMENTS1* (*PAP1*), which is a Myb-family master TF for induction of anthocyanin biosynthetic genes in response to several types of abiotic stresses (Liu et al., 2014; Shi and Xie, 2010; Shin et al., 2015). These defects were observed both at the ET and LT after salt stress except for *med9* which showed no defects in *PAP1* expression at either the ET or LT.

**Figure 7.**
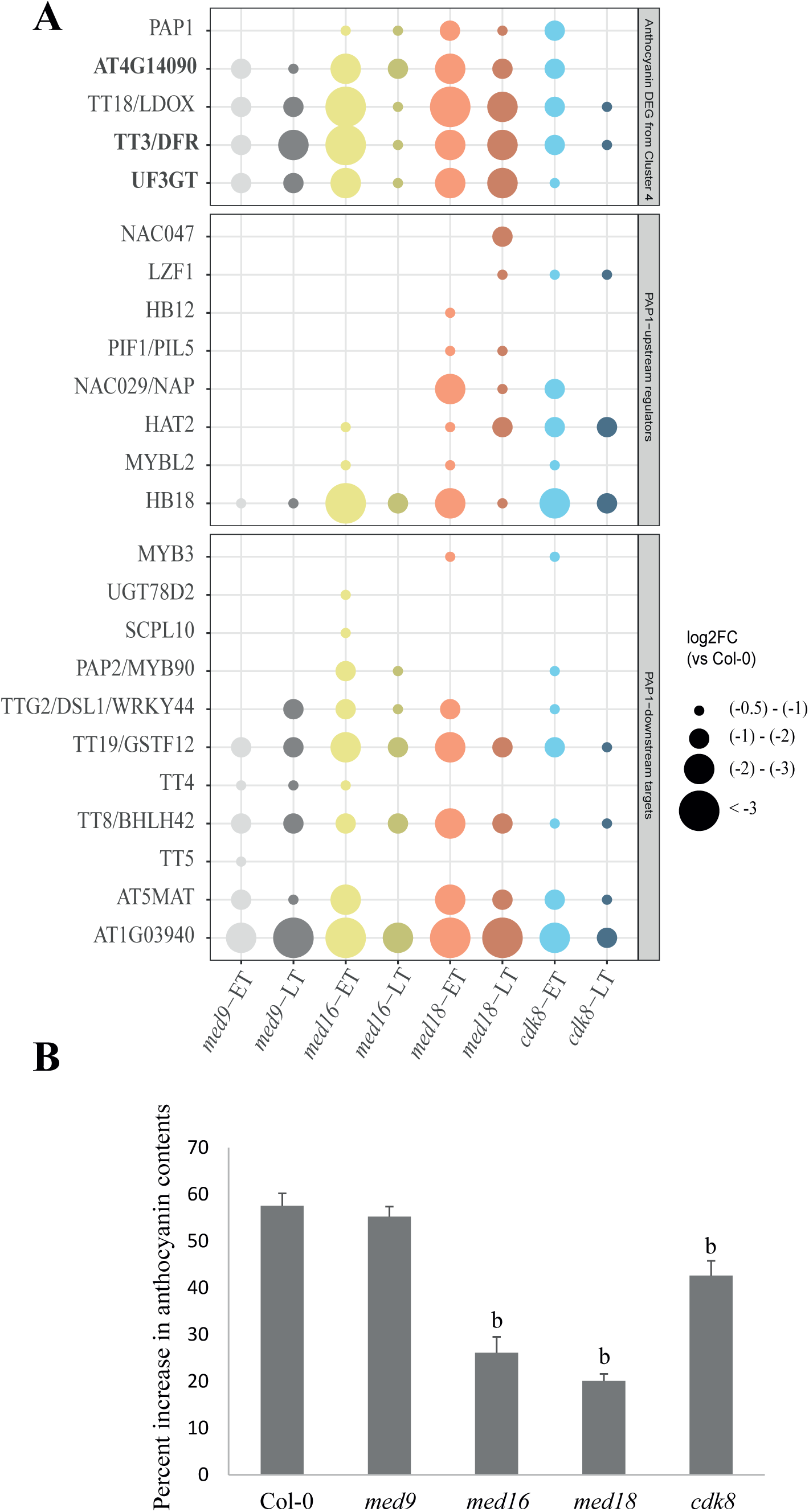
Mis-regulated DEGs encoding proteins involved in anthocyanin biosynthesis for *med9*, *med16*, *med18* and *cdk8*. **(A)** Fractions of misregulated DEGs in mutants at the ET and LT from the GO category “Anthocyanin−containing compound biosynthetic process” (GO:0009718) (upper panel), PAP1-upstream regulators (middle panel) and PAP1-downstream targets (lower panel). Genes in bold type in the upper panel are also PAP1-downstream targets. Dot size represents the ratios between the log2-fold change in expression at the ET or LT relative to control in mutants relative to Col-0. **(B)** Measurement of anthocyanin contents in *med9*, *med16*, *med18* and *cdk8* when irrigated with 300 mM salt solution for two weeks. Leaves were taken from 5-week-old plants irrigated with (300 mM) NaCl solution for15 days. Percent increase in anthocyanin contents of leaves was quantified. Three independent experiments were performed using 4 plants per experiment. Error bars indicate ± SD (n = 3). Statistical analyses were performed comparing Col-0 with each mutant separately. p≤0.05=a, p≤0.01=b (Student’s t-test).

The Plant Transcription Factor Database (Plant TFDB, v5.0) was used to identify eight TFBSs in the *PAP1* promoter which are specific for up-stream regulators of *PAP1* expression. Several of these TFs were mis-regulated in the Mediator mutants at the ET after salt stress (Fig. 7A middle panel; Table S24). *HB18* was identified as an NR gene in all four mutants, *HAT2* and *MYBL2* in *med16*, *med18* and *cdk8*, *NAP/NAC29* in *med18* and *cdk8*, *PIL5* and *HB-12* only in *med18*, and *LZF1* only in *cdk8*. The only TF-encoding gene that showed the same induction as Col-0 at the ET in response to salt stress was *NAC047*. However, *NAC047* showed reduced induction in *med18* at the LT. Plant TFDB was also used to identify fourteen downstream target genes for PAP1. Most of them were mis-regulated in all four Mediator mutants at both the ET and LT (Fig. 7A lower panel; Table S24). Interestingly, PAP1 has been shown to form a MBWW complex with other TFs (TTG1, TTG2 and TT8) to control pigment production in an auxin concentration-dependent manner (Cappellini et al., 2021; Shin et al., 2015). We found that expression of *TT8* was mis-regulated in all four mutants and *TTG2* in *med16*, *med18* and *cdk8*.

To connect our transcriptomic data to physiological effects, we quantified the anthocyanin levels in our mutants and Col-0 in response to salt stress. The anthocyanin contents were significantly reduced in *med16*, *med18*, and *cdk8* compared to Col-0 (Fig. 7B). Furthermore, both the anthocyanin levels (Fig. S6C), and the expression levels of *PAP1* (Fig. S7) were, at least partially, restored to the levels of Col-0 in the complemented lines. Our results show defects in induction of genes encoding proteins involved in anthocyanin metabolism, upstream regulation of *PAP1* expression and *PAP1* downstream targets. This was most pronounced in *med16* and *med18* and less in *med9* and *cdk8*. This is in line with our quantification of anthocyanin levels in Col-0 and the Mediator mutants. The reduced accumulation of anthocyanin content therefore provides one explanation for the salt sensitive phenotypes of *med16*, *med18* and *cdk8*.

### Genes involved in salt stress responses, ABA biosynthesis and ABA signalling in Cluster 2 show defects in med16 and med18 at the ET after salt stress

The results presented in Fig.s 1-3, indicate that *med16* and *med18* are the most salt stress sensitive of our Mediator mutants. This prompted us to perform a more detailed analysis of expression of genes involved in salt stress in these two mutants. We focused on NR genes in Cluster 2 since it includes genes belonging to GO categories related to salt stress (GO:0009651) and responses to ABA (GO:0009737) (Fig. 5D). The importance of ABA in salt stress responses is well documented and several of the NR genes for *med16* and *med18* in Cluster 2 are shared between these two GO categories. These are typically multi-task genes and have been shown to be important for responses to water deprivation, osmotic stress, signal transduction, membrane channels and lipid metabolism. A similar requirement of Med16 for expression of ABA-responsive genes was recently reported by others (Guo et al., 2021b; Lee et al., 2021)

Initially, we identified Cluster 2 genes which are NR in *med16*, *med18* or both, which encode proteins involved in signal transduction and are members of GO categories related to salt stress and/or responses to ABA (Fig. 8A). Mitogen-activated protein kinases (MAPK) are involved in pathways that transmit environmental cues to the transcription machinery, and they have been shown to be involved in salt stress responses (Sinha et al., 2011; Ulm et al., 2002). We found that expression of *MAP KINASE KINASE 9* (*MKK9*; *AT1G73500*) and *MAPKKK18* (*AT1G05100*) was significantly induced at the ET in Col-0 but showed a significantly lower induction in both *med16* and *med18* (Table S16, Cluster 2). For *med18*, we also identified *CALCINEURIN B-LIKE PROTEIN 1* (*CBL1*; *AT4G17615*), and *CALCIUM-DEPENDENT PROTEIN KINASE 32* (*CPK32*; *AT3G57530*) as significantly less induced at ET relative to Col-0 (Table S16, Cluster 2). Both *CPK32* and *CBL1* have previously been implicated in responses to both salt stress and ABA (Cheong et al., 2003; Li et al., 2017).

**Figure 8.**
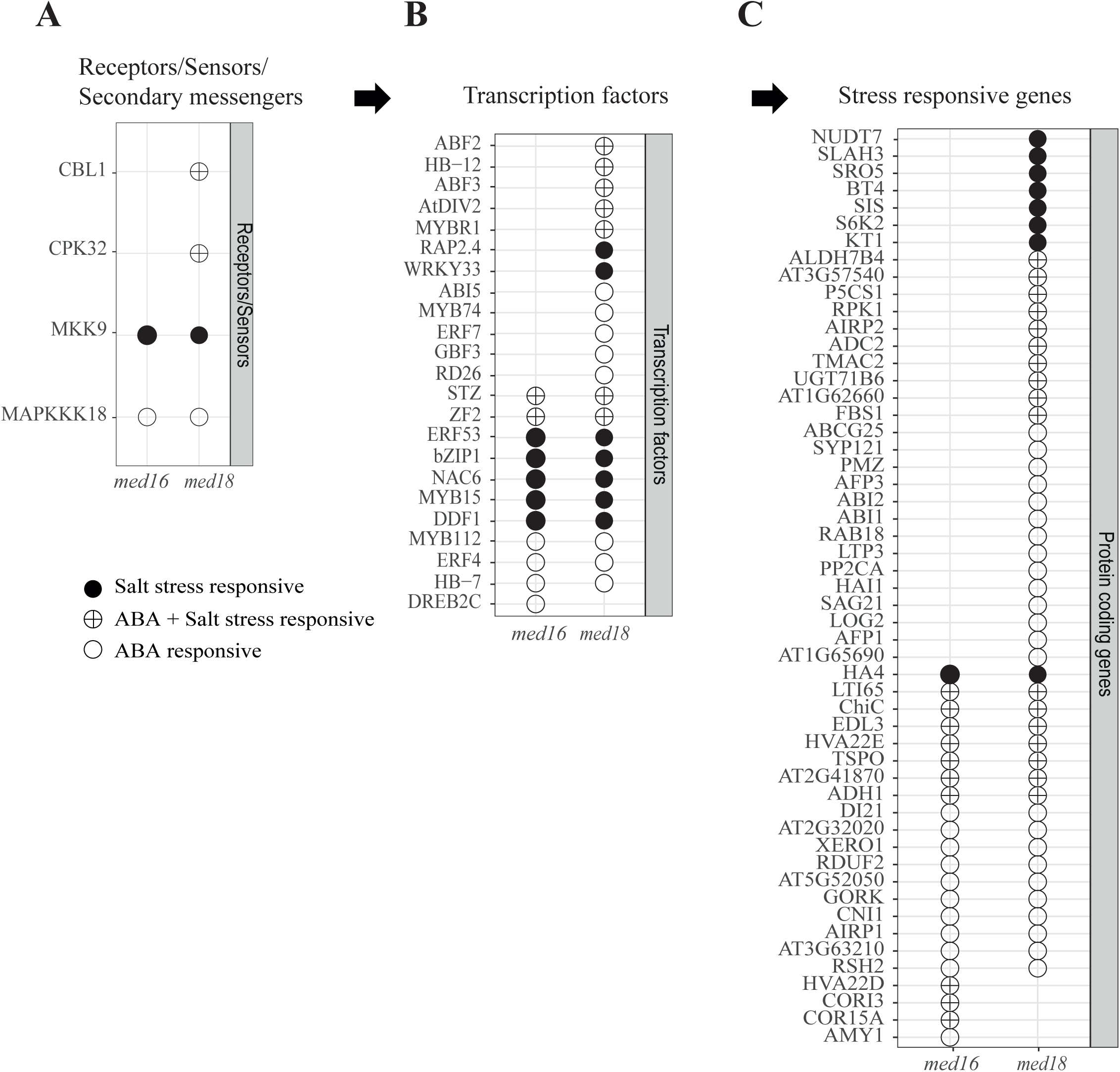
NR genes involved in salt stress responses and ABA biosynthesis and signalling in Cluster 2 of *med16* and *med18*. NR genes were obtained from the Cluster 2 GO categories “Response to salt stress (GO:0009651)”, and “Response to abscisic acid (GO:0009737), described in Figure 5D. The genes were separated into Sensors/Receptors/ Secondary messengers (A), Transcription factors (B), and Target genes (C) and analysed for their induction at the ET after salt stress in *med16* and *med18* relative to Col-0.

To identify potential transcriptional defects that could explain the sensitivity to salt stress in *med16* and *med18*, we next identified TFs that are NRs in these mutants and are members of the “salt stress response” or “ABA response” GO categories (Fig. 8B). For *med16*, we identified one unique TF (*DREB2C*; *AT2G40340*) which is a member of the ABA response GO category, as an NR gene. In *med18* we identified a set of TFs that are encoded by NR genes uniquely in *med18* and are members of either the salt stress response GO category (*RAP2.4*, *WRKY33*), the ABA stress response GO category (*ABI5*, *MYB74*, *ERF7*, *GBF3*, and *RD26*), or both (*ABF2*, *HB-12*, *ABF3*, *DIV2*, and *MYBR1*). Interestingly, Med18 association with the *HOOKLESS1* (*HLS1*) histone acetyltransferase is require for acetylation and transcriptional activation of the *ABI5* and *WRKY33* promoters (Liao et al., 2016). Finally, both *med16* and *med18* showed defects in induction of the salt stress response genes *ERF53*, *bZIP1*, *NAC6*, *MYB15*, and *DDF1* genes as well as the salt stress and ABA-response genes *STZ* and *ZF2* (Fig. 8B).

Finally, we identified a large set of 53 target genes in Cluster 2 which belong to the salt response and/or ABA response GO categories of NR genes in *med16* and/or *med18* (Fig. 8C). Of these, four were unique to *med16*, 31 were unique to *med18* and eighteen were shared by both mutants. RT-qPCR experiments showed that the expression levels for three of these target genes were restored in the *med16-*C and *med18-*C complemented lines (Fig. S7). These results emphasize that responses to salt stress and ABA are strongly connected and provides an explanation to the phenotypes we have observed for *med16* and *med18*.

## Discussion

We selected four subunits (Med9, Med16, Med18, and Cdk8), representing each of the four Mediator modules of *Arabidopsis thaliana*, to study their function in response to salt stress. RNA-Seq experiments on Col-0 (WT) and the four mutant lines from samples collected prior to and at two time points following application of salt stress showed that global transcriptional responses to salt stress in Col-0 at the ET were consistent with a model for a fast transcriptional response to salinity. We detected rapid closure of stomata, activation of chitin response pathways, and reduced auxin signalling; responses that protect plants from salinity by reducing water loss (Ribba et al., 2020; Munns and Tester, 2008; Brotman et al., 2012). The early response was followed by sustained induction of transcripts for protein modification and leaf senescence, while transcripts for primary/secondary metabolism and bioenergetics were repressed as Col-0 plants adjust to suboptimal conditions. Similar delayed responses to different types of stress have been previously reported confirming that our experimental conditions were appropriate (Zhu, 2002).

Direct effects on changes in expression of salt stress target genes are best revealed at the ET, while changes in gene expression at the LT might result from indirect effects. We therefore identified TFBSs, and their corresponding TFs enriched in promoters of DEGs in Col-0 at the ET. For induced DEGs, the ERF-, WRKY-, bHLH-, Myb-, bZIP, and NAC-families were most common. These families have been implicated in different types of abiotic stress responses in Arabidopsis (Banerjee and Roychoudhury, 2015, 2017; Guo et al., 2021a; Shao et al., 2015; Wang et al., 2021; Xie et al., 2019). For the 50 most highly enriched individual TFs, we found that 75% of them belonged to the bZIP and bHLH-families. Nine of the ten most highly enriched TFs were bZIP-type TFs and they have all been connected to responses to the stress-related ABA hormone (Bensmihen et al., 2002; Choi et al., 2000; Hsieh et al., 2012; Yang et al., 2018, 2016). This was further supported by GO analyses showing that the category “Response to abscisic acid (GO:0009737)” was one of the most common among the ET DEGs. Another interesting finding among the 50 most enriched individual TFs was a group of six TFs belonging to the CCA/LHY/RVE family, which are most known for their function in control of the Arabidopsis circadian clock (Gray et al., 2017). However, a recent report suggested that Rve4 and Rve8 might function in response to heat stress (Li et al., 2019). Our results support and extend this finding and suggest that the entire CCA/LHY/RVE family might have a more general function, at least in response to salt stress.

The most common TF-families in promoters of repressed DEGs at the ET belonged to the ERF-, Myb-, bZIP-, HD-ZIP-, MIKC/MADS-, and Dof-families. The three latter were unique for repressed DEGs at the ET. The clearest differences between induced and repressed DEGs at the ET were that TFBSs for MIKC/MADS and HD-ZIP TFs were more than twice as common among the repressed DEG promoters compared to the induced DEG promoters, while TFBSs for WRKY and NAC TFs were 20- and 4-fold more common in the induced DEG promoters compared to the repressed DEG promoters. Of the fifty most enriched individual TFs, the most common belonged to the TCP-, Myb- and AGL-families. Interestingly, TCPs are most known for their involvement in regulation of multiple aspects of plant development, but recent reports show that they are also important to balance development and stress defense (Li, 2015). TCP5, TCP13, and TCP17, have been identified as positive regulators in plant response to heat stress (Han et al., 2019). They function together with the transcriptional activator PIF4, which is a central integrator of heat stress response where TCP5, TCP13, and TCP17 together with PIF4 activate a set of target genes in response to heat of which the most important are *PRE1* and *YUC8*. Our results confirm these functions but suggest that the effect is the opposite in salt stress since we found that expression of both TCPs and PIF4 is repressed in response to salt stress. Accordingly, our data show repression of the TCP/PIF4 target genes *PRE1* (6-fold at ET/564-fold at LT) and *YUC8* (3.5-fold at ET/11-fold at LT).

To find out if subunits from different Mediator modules are involved in specific regulatory transcription programs, we analysed coordinated gene expression responses to salt stress by means of heat maps and hierarchical clustering analysis of DEGs in Col-0 at the ET. GO and TFBS analyses of DEGs in each co-expressed cluster revealed that they were enriched for genes connected to unique combinations of GOs, indicating that genes in each cluster are controlled by unique transcriptional programs. This was supported by analyses of enriched TFBSs in their promoters. At the level of TF-families, we could both identify families that were unique, *e.g.,* WRKYs for Cluster 2, Myb-related for Cluster 3, and TCPs, MIKC-MADSs, and MYBs for Cluster 5. However, we were surprised to find large overlaps between clusters. Most strikingly, the promoters of DEGs in Clusters 1 and 4, which include genes that respond in opposite ways to salt stress, where both enriched for the same TF-families (ERFs, bZIPs and bHLHs). In fact, the ERF-family was enriched in all clusters except Cluster 5. However, comparison at the level of specific, individual TFs, we found that the ERFs enriched in DEGs promoters of the repressed Cluster 1 differed markedly from the ERFs enriched in the induced Clusters 2-4. Apparently, individual members of the ERF-family can have opposite functions in response to stress.

Our phenotypic analyses revealed effects at both global and specific levels and showed that removal of single subunits from each module cause both common and unique defects in their salt stress response. For example, all four Mediator mutants showed a higher percentage of ion leakage compared to Col-0. In addition, *med9, med16* and *med18* displayed reduced primary root lengths and *med16*, *med18* and *cdk8* showed defects in anthocyanin metabolism. Finally, *med16* and *med18* showed reduced ability to protect their chlorophyll from damaging effects of salt stress by displaying more bleached leaves compared to Col-0. These phenotypic data were supported by gene expression defects in the mutants. At the global level, *med9* had the fewest number of mis-regulated genes relative to Col-0 in response to salt stress and consequently displayed the mildest phenotypic effects of the mutants. In contrast, *med18* displayed several severe phenotypes and accordingly showed the highest number of mis regulated genes compared to Col-0.

At a more detailed level, we found that NR genes in Cluster 3 for all mutants were enriched for the cell-cycle related GO categories “Microtubule−based movement” and “Regulation of cyclin−dependent protein serine/threonine kinase activity”. This was supported by our analysis of promoters in Cluster 3, which showed enrichment of a specific subset of MYB TFs called “MYB3R” in all mutants. The MYB3R family has been shown to bind to the MSA (M-specific activator) element of B-type cyclins to control their periodic promoter activation and to promote entry into mitosis (Haga et al., 2011; Ito et al., 2001). Dysfunctional regulation of cell-cycle regulated genes due to defects in function of MYB3R TFs in our four mutants might thus explain both their increased ion leakage and reduced primary root length phenotypes.

The observed phenotypic defects in anthocyanin metabolism and the enrichment of the GO category “Anthocyanin−containing compound biosynthetic process” among NR genes in Cluster 4 of NR genes in *med16*, *med18* and *cdk8* was intriguing, but we were unable to identify TFs that were significantly enriched in the NR gene promoters of Cluster 4 in any of our mutants. Our analyses of enriched TFBSs in promoters of NR genes in different cluster and mutants was designed to identify TFs that, for their function, are dependent of interaction with a specific Mediator subunit. However, the effect of Mediator mutations on expression of target genes can also result from changes in expression levels of the TFs themselves. Interestingly, we found that *PAP1*, a master TF for transcription activation of anthocyanin biosynthetic genes, was identified as an NR gene in Cluster 4 of *med16*, *med18* and *cdk8*. It is likely that mis-regulation of *PAP1* expression is a major cause of the observed defects in anthocyanin production in our mutants, since we also found that a majority of eight upstream TFs that might control *PAP1* expression and fourteen downstream *PAP1* target genes were NR genes in the mutants. We noticed that *med16* and *med18*, which displayed the largest defects in anthocyanin induction in response to salt stress, also showed the largest defects in expression of genes up- and downstream of *PAP1*.

Finally, we aimed to provide an explanation to the salt stress sensitivity of the most affected mutants, *med16* and *med18*. Our GO analysis revealed that NRs in Cluster 2 of *med16* and *med18* were enriched for genes encoding proteins involved in responses to salt stress and abscisic acid. By analysing the individual genes that contributed to these GOs among the NR genes in Cluster 2 of *med16* and *med18*, we found that they encode proteins that act at both the level of upstream signal transduction pathways, as transcriptional regulators, and as downstream target genes. There was a large overlap of NRs between *med16* and *med18* where nearly all NRs in *med16* were also identified as NRs in *med18*. However, we also identified a large set of NRs that were unique for *med18*, which might explain its more severe sensitivity to salt stress.

In summary, we show that deletions of subunits from each of the four Mediator modules affect the Arabidopsis transcriptional responses to salt stress at the global level, where specific mutants display both unique and overlapping phenotypes in response to salt stress. These phenotypes can be linked to the inability of certain TF-families to function in mutants lacking individual Mediator subunits. In some cases, we could identify individual TFs that depend on a specific Mediator subunit for a proper transcriptional response to salt stress. Focusing on genes in specific clusters that encode proteins connected to specific GO categories in salt stress response and which are NR in Mediator mutants, we identify genes involved in ABA and anthocyanin metabolism which indicates that genes belonging to these processes are dependent on specific Mediator subunits. The results presented here provide a starting point to study interactions between TFs and Mediator subunits to reveal how signals from different stress response pathways are integrated by Mediator subunits and subsequently encoded for a proper cellular salt stress response.

## Methods

### Plant materials

*Arabidopsis thaliana* ecotype Columbia (Col-0) was used as a control and genetic background for all lines in this study. Mutant lines of *med9* (SALK_029120), *med16* (SALK_048091), and *med18* (SALK_027178) are T-DNA-insert knockouts and were obtained from the Nottingham Arabidopsis Stock Centre (NASC; Nottingham, UK). The *cdk8* mutant line (GABI_564F11) used here has been described previously (Ng et al., 2013). All Mediator knock-out lines were verified as homozygous for the appropriate T-DNA insertion and transcript levels were quantified by RT-qPCR using a Roche 454 (Roche, Clifton NJ, USA) as described previously (Crawford et al., 2020).

### Growth conditions and salt treatment

Plants were grown in plates or on soil for physiological characterization under salt stress, either at the seedling stage or at the 5-week-old plants stage. For salt stress RNA-Seq experiments, plants were grown in a hydroponic system. Seeds were sterilized and subjected to cold treatment at 4℃ for 72 hours before being sowed either on plates, Eppendorf holder or in soil. Seeds were germinated at 22/18℃ (day/night) under short days (SD) (8h light: 16h dark) or long days (LD) (16h light: 8 h dark) in soil. When grown on plates or hydroponically, only SD conditions were used. Growth rooms were equipped for maintaining 65% air humidity under standard illumination (cool white light provided by fluorescent tubes (Osram Lumilux L18W 840 (Germany); peak wavelength 840 nm) at a photon flux density of 120 µE x m^-2^ x s^-1^. ½ x MS media (Duchefa Biochemie) was used as a basal medium with 0.5% sucrose for seedling stage experiments. pH for the medium was adjusted to 5.7, followed by the addition of 7 g/L phytoagar for solid medium only and subjected to autoclaving. For salt treatment, seedlings were grown for 5-7 days on MS media and then transferred to either MS media or MS media containing 0.2 M NaCl.

For the hydroponic system, a protocol was adopted from (Conn et al., 2013). In brief, seeds were sowed in punctured holes of 1.5 ml black Eppendorf caps filled with 250-300 µl of solidified germination media and then placed in floating racks filled with liquid germination media. Germination media in the floating rack was replaced with Basal nutrient media on the third day after germination. Seedlings were grown in floating racks for 3 weeks and then transferred to large, aerated containers. Each container contained 10 plants of each genotype and there were 3 containers each for control and salt-treated plants in each experiment. The experiment was replicated 4 times and samples were collected for RNA-Seq analysis.

For soil-grown plants, seeds were sowed directly into the soil. Five-week-old plants were irrigated with a final concentration of 0.3 M NaCl solution and were used for phenotyping and measurement of ion leakage after 1 week. Leaves of 5-week-old plants were incubated in 0.2 M NaCl solution for the indicated number of days and used to quantify the loss in chlorophyll content.

### Measurement of physiological parameters: root length, ion leakage rate, chlorophyll contents, and anthocyanin contents

Seven-day-old seedlings were transferred to either ½ x MS or ½ x MS + NaCl media. After one additional week, seedlings were examined for chlorosis/bleaching and bleached leaves were quantified. Measurements of the primary root lengths were performed two weeks after transfer from normal media. Images were taken and processed using the image J software to quantify the reduction in root length.

Mature plants (five-weeks-old) were irrigated with either water or salt solution and used for measurement of ion leakage rate/electrolyte leakage using the method described in (Julkowska et al., 2016). In brief, two leaf disks were taken from mature leaves at 72 hours after initiation of the salt stress treatment. Leaf discs were washed with MQ water for 30 minutes by shaking under light and then transferred to 0.1 % w/w) Silwet solution. The samples were vacuum infiltrated twice for 2 minutes each and then incubated for 1 hour at an orbital shake. Ion leakage was then determined by measuring the conductivity of the solutions using an electrical conductivity meter (CDM210, conductivity meter, Meterlab). Total ion leakage was measured after boiling the samples. Percent increase in ion leakage after salt treatment was quantified for Col-0 and mutant lines.

Chlorophyll contents were determined as described (Hiscox and Israelstam, 1980). Detached leaves were harvested from 5-weeks-old plants and incubated in 0.2 M salt solution for either 2 days (*med16*) or 4 days (*med18*). 50 mg of leaves were used and incubated in 10 ml DMSO at 65 ℃ for 5 hours. Samples were then cooled to room temperature. Absorbances of samples were recorded at 663 and 645 nm. Chlorophyll contents were calculated using the following equations as described in (Arnon, 1949); For chlorophyll a (mg g^-1^ FW) = [(12.7 x A_663_) - (2.69 x A_645_)] x V/1000 x W, For chlorophyll b (mg g^-1^ FW) = [(22.9 x A_645_) - (4.68 x A_663_)] x V/1000 x W, Total chlorophyll a + b (mg g^-1^ FW) = [(8.02 x A_663_) + (20.2 x A_645_)] x V/1000 x W, where V is the final volume of DMSO solution in ml and W is the fresh weight of plants material.

Anthocyanin contents were accessed after salt treatment using the method described in (Zhao et al., 2018). Plants were irrigated with water or a 0.3 M NaCl solution and samples for anthocyanin contents assessment were collected after two weeks. 50 mg of leaf samples were collected and incubated in a 600 µl methanol solution containing 1% HCl (v/v). Samples were incubated at 4 ℃ overnight with gentle shaking. 400 µl of water and 400 µl of chloroform were added and samples were mixed by the vortex. Samples were then centrifuged at 12,000 x g for 2 minutes and the resulting supernatants were used to measure the absorbance at 530 and 657 nm. Anthocyanin contents were calculated using the formula A_530_ - 0.25 A_657_. All experiments were conducted in triplicates and student’s t-test was applied for statistical analysis.

### RNA isolation and RT-qPCR

Total RNA was extracted from rosette of plants gown under control or salt stressed conditions. 100 mg of fresh material was grounded in liquid nitrogen and RNA was extracted using the E.Z.N.A Plant RNA kit (Omega Bio-tek, Norcross, USA). Samples were processed to remove DNA contamination following the protocol provided Turbo DNA-free DNase I (Ambion, Foster City, USA) kit. RNA samples were then quantified and checked for RNA integrity using an Agilent BioAnalyzer 2100 with RNA Nano 6000 kit (Agilent Technologies, Santa Clara, USA), and only samples with an RNA integrity number ≥8 were sent for RNA-Seq. Similarly prepared samples were used for RT-qPCR analysis.

1 µg of total RNA was reverse transcribed using iScript reverse transcription supermix (Biorad, Solna, Sweden). RT-qPCR was performed using a LightCycler 96 and the SYBR green master mix (Roche, Solna, Sweden). Gene expression levels were normalized to the AT5G09810 and AT5G25760 reference genes and displayed in relative units. Two biological and three technical repetitions were performed for each sample. The sequences of primers (Mahmood et al., 2016; Ryu et al., 2010) used for the RT-qPCR primers are shown in Table S1.

### Construction of complemented lines

An 8,117 bp fragment of *MED16* and a 2,068 bp fragment of *MED18* including ∼1,000 kb promoter sequences and the coding regions were produced using PCR and primers shown in Table S1.

The synthesized DNA fragments were incorporated into the gateway vector pDonor221 by homologues recombination and verified by sequencing before re-ligated into the target plasmid pGWB513-3xHA. The recombinant binary vectors were transfected into *Agrobacterium* GV3101 using electroporation. The *Agrobacteria* strains were then used to transform the *med16* and *med18* T-DNA lines using the floral dip transformation procedure. The primary transformants were selected on ½ x MS plates containing 30 µg/ml hygromycin and 150 µg/ml timentin. The positive transgenic plants were identified using RT-PCR and primers specific for the genomic gene of interest fused to the HA-tag. Homozygous lines were obtained by propagating the transgenic plants for another two generations.

### Pre-processing of RNA-Seq data and Identification of Differentially Expressed Genes

RNA-Seq library construction was completed by NGI Uppsala as described earlier (Crawford et al., 2020). Raw RNA-Seq data was pre-processed by NGI Uppsala, according to best practice using TrimGalore and FastQC. Read counts were obtained using the kallisto R package (v0.43.0) (Bray et al., 2016) and mapped to the Araport11 Arabidopsis Col-0 reference genome annotation (Cheng et al., 2017). The code was executed as follows: kallisto quant -i kallisto/Araport11_all.201606.cdna.inx -b 100 -o kallisto/P6960_144_R1_trimmed --rf-stranded - t 8 --rf-stranded trim_galore/P6960_144_R1_trimmed.fq.gz -l 250 -s 50. Uniquely-mapping transcripts were counted and expressed as transcripts per million. The kallisto abundance values were imported into R (v3.3.2; R Core Team 2015) with the help of the Bioconductor (v3.3) (Gentleman et al., 2004). tximport package (v.1.2.0) (Soneson et al., 2016). Beginning with a detected population of around 26,000 out of 32,834 annotated features such as genes and ncRNAs, lowly expressed transcripts (less than one transcript, in no more than 2 of 4 replicates in any condition) were filtered out using a custom feature-select script to produce 25,914 genes. Direct comparison of global transcriptomes was achieved with Principal Component Analysis (PCA), performed using the R core package prcomp, or by creating heatmaps using custom R scripts. To normalize the data and enable better comparison between highly and lowly expressed genes, a variance-stabilizing transformation was applied on the raw data using the Bioconductor DESeq2 package (v1.14.1) (Love et al., 2014). Clustering and scaling of variance-stabilized data was performed using Ward’s method of Pearson correlation. Heatmaps and bubble plots were generated in R using the ggplot2 package (v2.3.0). Heatmaps of NR genes were generated using Complex Heatmap package (v2.7.9.1000) (Gu et al., 2016).

### Gene ontology analysis for enriched functional categories

DEGs identified for each genotype were analysed for functional categories enrichment analysis using the Gene ontology (GO) databases. GOs were assessed using a custom R script developed in our lab. The database used in the script for the analysis was obtained from TAIR (https://www.arabidopsis.org/download/index-auto.jsp?dir=%2Fdownload_files%2FGO_and_PO_Annotations%2FGene_Ontology_Annotations; database file: ATH_GO_GOSLIM, 2019-07-11). Enriched functional categories with a Bonferroni corrected *p*-value < 0.05 were selected and represented in terms the ratio between the number of genes observed for a specific category in our study, and the number of genes that has been allocated to that specific functional category in the entire background dataset of *A. thaliana*. The number of enriched GO functional categories was reduced using REVIGO (http://revigo.irb.hr/) by selecting the option to remove obsolete GO terms and a database that was set to “Arabidopsis thaliana (3702)”.

### Transcription factor binding site enrichment analysis

Enrichment of transcription factor (TF) binding sites in the four stress regulons was analysed using TF2Network (Kulkarni et al., 2018). A p-value of < 0.05 was set as the threshold for significance. The output file from TF2Network generally contained multiple binding sites for the same TF. For some analyses data were collapsed to the level of TF-families, which we defined according to the Arabidopsis specific file from PlantTFDB 5.0 (Tian et al., 2020a). To test for enrichment of any of TF binding sites (TFBSs) in the subgroup of the stress regulon genes which did not respond to stress in the mutants (‘non-responsive’), the number of targets (Positional weight matrix (PWM)) in the subgroup ‘non-responsive’ for each mutant and cluster were compared to the total number of targets in the corresponding cluster of Col-0. Hypergeometric distribution was used to test for enrichment of specific TFBSs in subgroups of genes. P-values were calculated using the phyper function in R and a Bonferroni corrected p-value of 0.05 was used as a significance threshold.

### Accession Numbers and Data Access

The sequencing data has been deposited at the European Nucleotide Archive (ENA, ) under accession number PRJEB33339.

## Acknowledgements

We are indebted to the service, support and training provided by the UPSC bioinformatics platform, especially the talents of Dr. Nicolas Delhomme. We thank Dr. Inge de Clercq (La Trobe University, Australia) for seeds of the *cdk8* mutant. This work was supported by the Knut and Alice Wallenberg Foundation [2015-0056 to S.B and Å.S.], by the Swedish Foundation for Strategic Research [SB16-0089 to Å.S. and S.B.] and by the Swedish Research Council [2016-03943 to S.B., and 2016-04319 to Å.S].

## Data Availability

The sequencing data has been deposited at the European Nucleotide Archive (ENA, www.ebi.ac.uk/ena) under accession number PRJEB33339.

## Author contributions

F.K. and J.B. designed the research, performed salt stress experiments, analysed the RNA-seq results and performed phenotypic experiments, A.V. Performed all bioinformatic analyses of RNA-seq data together with M.R. and produced figures related to RNA-seq. T.C and N.L participated in salt stress experiments. Å.S. and S.B. designed the research and wrote the paper.

## Statement

None of the authors have any competing interests.

